# Axon-like protrusions promote small cell lung cancer migration and metastasis

**DOI:** 10.1101/726026

**Authors:** Dian Yang, Fangfei Qu, Hongchen Cai, Chen-Hua Chuang, Jing Shan Lim, Nadine Jahchan, Barbara M. Grüner, Christina Kong, Madeleine J. Oudin, Monte M. Winslow, Julien Sage

## Abstract

Metastasis is the main cause of death in cancer patients but remains a poorly understood process. Small cell lung cancer (SCLC) is one of the most lethal and most metastatic types of human cancer. SCLC cells normally express neuroendocrine and neuronal gene programs but accumulating evidence indicates that these cancer cells become relatively more neuronal and less neuroendocrine as they gain the ability to metastasize. Here we show that mouse and human SCLC cells in culture and *in vivo* can grow cellular protrusions that resemble axons. The formation of these protrusions is controlled by multiple neuronal factors implicated in axonogenesis, axon guidance, and neuroblast migration. Disruption of these axon-like protrusions impairs cell migration in culture and inhibits metastatic ability *in vivo*. The co-option of developmental neuronal programs is a novel molecular and cellular mechanism that contributes to the high metastatic ability of SCLC.

## INTRODUCTION

Metastases are a major cause of cancer-related morbidity and mortality. By the time cancer cells leave their primary site and spread to distant sites, they have acquired the ability to migrate and invade, as well as characteristics that enable them to survive and proliferate within new microenvironments. These phenotypes are likely driven by changes in gene expression and epigenetic programs that allow cancer cells to overcome the many hurdles that normally constrain the metastatic process. Despite recent advances, our understanding of the principles and mechanisms underlying metastasis remains incomplete, including how changes in molecular programs can translate into selective advantages that enable cancer cells to spread to other organs (Fidler, 2003, Lambert *et al.*, 2017, Obenauf and Massague, 2015).

Small cell lung carcinoma (SCLC) is a high-grade neuroendocrine cancer that accounts for ∼15% of all lung cancers and causes over 200,000 deaths worldwide each year (Sabari *et al.*, 2017). The ability of SCLC cells to leave the primary tumor and establish inoperable metastases is a major cause of death and a serious impediment to successful therapy (Farago and Keane, 2018, van Meerbeeck *et al.*, 2011). SCLC is one of the most metastatic human cancers, with over 60% of SCLC patients presenting with disseminated disease at the time of diagnosis, often including liver, bone, brain, and secondary lung metastases (Nakazawa *et al.*, 2012, Riihimaki *et al.*, 2014). Molecular analyses to understand metastatic progression of human cancer are often limited by difficulties in accessing tumor samples at defined stages. This problem is especially true for SCLC, since patients with metastatic disease rarely undergo surgery (Barnes *et al.*, 2017). Genetically engineered mouse models of human SCLC recapitulate the genetics, histology, therapeutic response, and highly metastatic nature of the human disease (Gazdar *et al.*, 2015, Kwon and Berns, 2013, Rudin *et al.*, 2019). These genetically engineered mouse models recapitulate cancer progression in a controlled manner and allow for the isolation of primary tumors and metastases directly from their native microenvironment. Recently, we and others have used mouse models to uncover gene expression programs that are enriched in SCLC metastases (Denny *et al.*, 2016, Semenova *et al.*, 2016, Wu *et al.*, 2016, Yang *et al.*, 2018). While SCLC cells display features of neuroendocrine cells, the gene expression programs in metastatic SCLC include not only genes normally expressed in pulmonary neuroendocrine cells but also those expressed in neurons (Carney *et al.*, 1982, Cutz, 1982, Broers *et al.*, 1987, Anderson *et al.*, 1988). Higher levels of the neuronal markers such as NSE (neuron-specific enolase) correlate with shorter survival and more metastatic disease in SCLC patients (Carney *et al.*, 1982, Dong *et al.*, 2019, van Zandwijk *et al.*, 1992). Broad neuronal gene expression programs are enriched in metastases from mouse models of SCLC, however, whether SCLC cells actually gain neuronal characteristics and whether neuronal features are key regulators of metastatic ability has not been previously characterized (Denny *et al.*, 2016, Wu *et al.*, 2016, Yang *et al.*, 2018, Böttger *et al.*, 2019).

Here we find that the metastatic state of SCLC is linked to the growth of protrusions that resemble axons. These axon-like growths increase the ability of SCLC cells to migrate and metastasize, thus representing a cellular mechanism that enhances the metastatic ability of SCLC cells that have transitioned to a more neuronal cell state.

## RESULTS

### SCLC cells can form long cellular protrusions in culture and *in vivo*

To investigate SCLC migration, we developed an assay in which SCLC cells, which classically grow in culture as floating spheres or aggregates, are grown as a monolayer under Matrigel ((Denny *et al.*, 2016) and Methods). Unexpectedly, we noticed that cells from some SCLC cell lines (N2N1G, 16T, 6PF) derived from the *Rb* ^*f/f*^;*p53*^*f/f*^ (*DKO*) and *Rb* ^*f/f*^;*p53* ^*f/f*^;*p130* ^*f/f*^ (*TKO*) genetically engineered mouse models form long cellular protrusions into cell-free spaces (Figure 1A-B). To determine whether these structures specifically project into cell-free areas or they also exist within monolayers, we cultured a minor fraction of fluorescently-labeled, GFP^positive^ SCLC cells with control SCLC cells. We found that SCLC cells also form protrusions when they are in close contact with surrounding cancer cells (Figure S1A). Similar mixing experiments performed in subcutaneous allografts also documented the growth of protrusions by SCLC cells *in vivo* (Figure 1C-D). Finally, similar structures also extend from SCLC micro-metastases in the liver in the autochthonous *TKO* mouse model and after intravenous transplantations of SCLC cells (Figure S1B-C).

**Figure 1:**
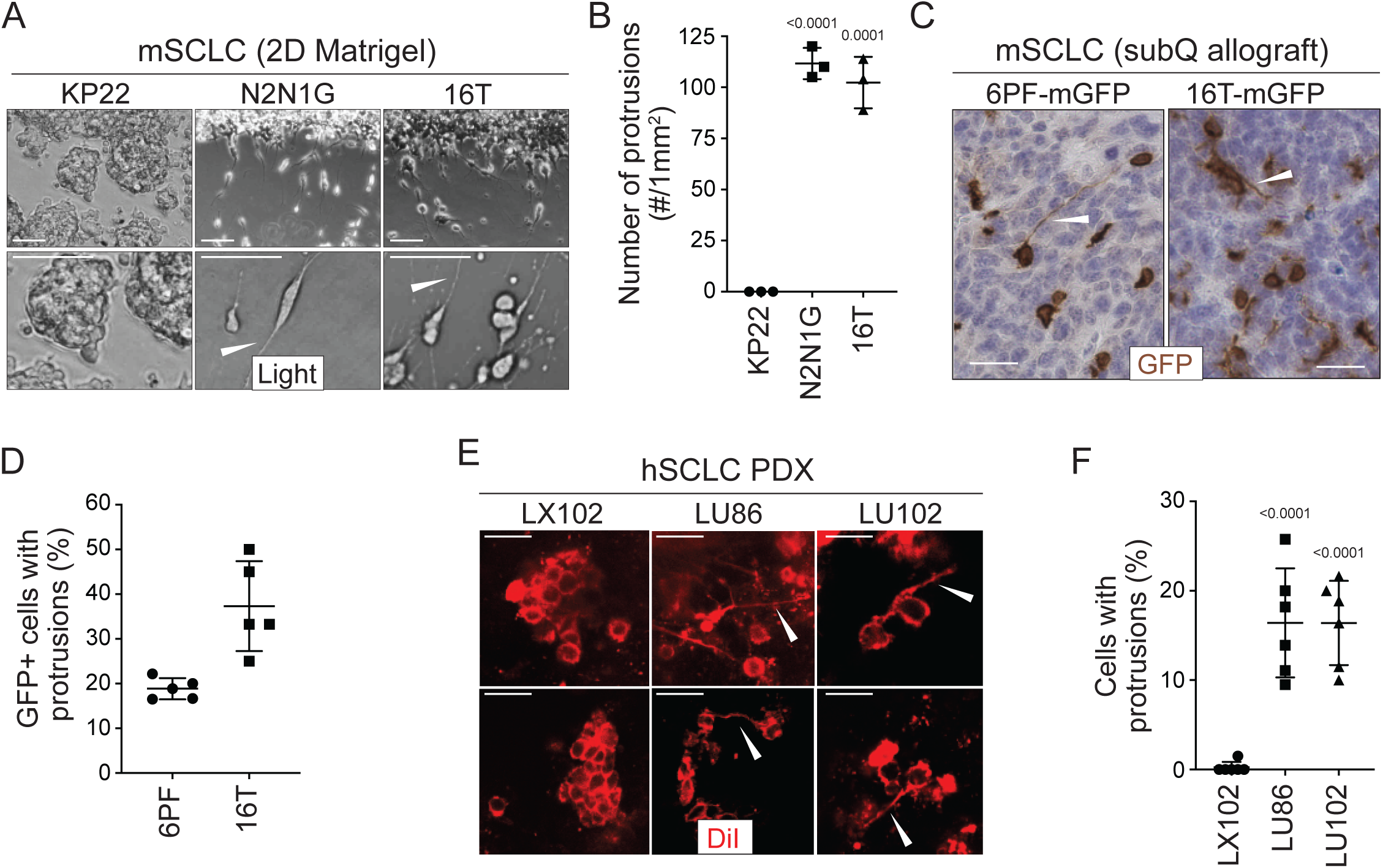
SCLC cells grow protrusions in culture and *in vivo*. A. Representative bright field images of three mouse SCLC (mSCLC) cell lines (KP22, N2N1G, and 16T). Cells extend protrusions into a cell-free scratch generated in monolayer cultures. Protrusions are shown with white arrowheads. Scale bars, 100 μm. N=3 replicates. B. Quantification of the number of protrusions that form from each mSCLC cell line as cultured in (A). Each symbol corresponds to the average of two technical replicates of an independent experiment. Mean +/− s.d. is shown, unpaired t-test. C. Representative images of mSCLC cells (6PF and 16T) growing as subcutaneous tumors. At the time of injection, 10% SCLC cells stably expressing membrane-GFP (mGFP) were mixed with 90% GFP-negative SCLC cells. Immunostaining for GFP generates a brown signal. Examples of protrusions are shown with white arrowheads. Hematoxylin (blue) stains the nuclei of the cells. (N=5/allograft, from one biological replicate). Scale bar, 20 μm. D. Quantification of (C). Each symbol represents an allograft tumor (N=4/allograft, from one biological replicate). Mean +/− s.d. is shown. E. Representative images of human SCLC (hSCLC) patient derived xenografts growing subcutaneously (LX102, LU86, and LU102 models). Tumors were injected with the red fluorescent tracer DiI. Protrusions are shown with white arrowheads. Scale bar, 20 μm. F. Quantification of (E). Each symbol represents a xenograft tumor (N=6/xenograft, from one biological replicate). Mean +/− s.d. is shown, unpaired t-test.

Human SCLC patient-derived xenografts (PDXs) recapitulate many important features of the human disease (e.g. (Gardner *et al.*, 2017, Saunders *et al.*, 2015)). To label rare cancer cells within human SCLC PDXs and identify whether they had protrusions in unperturbed tumors, we used DiI tracing. DiI is a lipophilic dye that diffuses within cell membranes and has been widely employed to label projections from individual neurons (Heilingoetter and Jensen, 2016, Mufson *et al.*, 1990). Protrusions from SCLC cells were easily identifiable in two out of three PDX models (Figure 1E-F). In the 2D monolayer culture system, not all human SCLC cell lines formed protrusions, but NCI-H446 cells formed long protrusions into cell-free areas analogous to those that formed in human SCLC PDX (Figure S1D-E). NCI-H446 cells also formed protrusions when grown as xenografts (Figure S1F).

These observations indicated that at least a subset of SCLC cells, which are often described as being “small round blue” cells, can develop long cellular protrusions. We next sought to investigate the nature of these protrusions and uncover their possible role in metastatic SCLC.

### SCLC protrusions resemble axons and SCLC cells with protrusions migrate similar to neuroblasts

SCLC cells express typical neuroendocrine genes but also neural and neuronal genes (Carney *et al.*, 1982, Cutz, 1982). This observation led us to investigate whether the protrusions were similar to neuronal axons or dendrites. We identified a list of 70 genes classically associated in the scientific literature with axonogenesis and axon guidance, and found that many of these genes are expressed in at least subsets of primary human SCLCs (George *et al.*, 2015) (Table S1). Thus, the gene expression programs controlling axonal growth in neuronal cells are also present in SCLC cells. We previously performed gene expression analyses on purified cancer cell from primary tumors and metastases from two mouse models of SCLC (Denny *et al.*, 2016, Yang *et al.*, 2018). In these studies, we found a general increase in the expression of neuronal gene expression programs during tumor progression. Indeed, almost all (69/70) of the selected candidate genes were expressed in metastatic SCLCs, indicating that murine SCLC tumors and cell lines derived from these tumors represent a tractable system with which to investigate neuronal programs in SCLC (Table S2). Pathway and process enrichment analysis on these 69 genes confirmed their connection with axon guidance, neuron migration, and nervous system development (Table S3).

To further investigate the nature of these SCLC protrusions, we assessed their expression of canonical axonal and dendritic proteins. The protrusions that form from murine and human SCLC cell lines were uniformly positive for the expression of neuron-specific class III beta-tubulin (Tuj1). More importantly, these protrusions were positive for the axonal marker TAU while expression of the dendritic marker MAP2 was undetectable (Figure 2A-B and Figure S2A-C). Tuj1^positive^, TAU^positive^ protrusions were also observed *in vivo* emanating from SCLC cells in the liver of *TKO* mice (Figure S3A). Furthermore, ∼37% (29/79) of human primary SCLC tumors stained moderately or strongly positive for TAU (Figure S3B). Together, these observations showed that SCLC tumors express axonal markers in different contexts and suggested that the protrusions observed on SCLC cells are axon-like.

**Figure 2:**
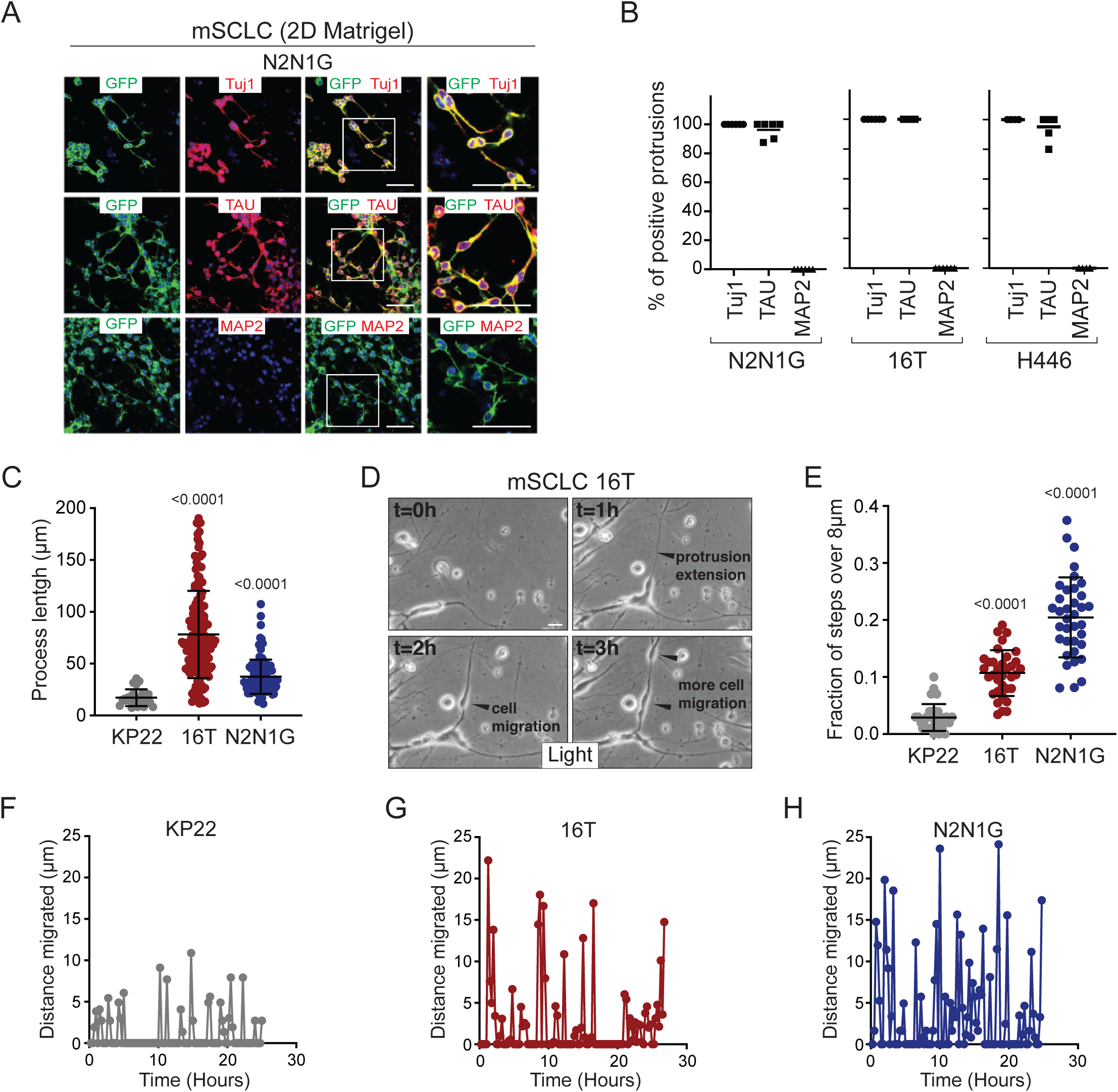
SCLC cells with protrusions migrate in a saltatory fashion similar to neuroblasts. A. Representative immunofluorescence images of N2N1G mSCLC cells expressing membrane-GFP (GFP, green) and stained (red) for expression of the neuronal marker Tuj1, the axonal marker TAU, or the dentritic marker MAP2. DAPI marks the nucleus of cells in blue. Scale bars, 50 μm. B. Quantification of (A) for two mouse SCLC cell lines (16T, N2N1G) and one human SCLC cell line (H446). Images for 16T and H446 are shown in Figure S2B-C. N=5/cell line. The bar is the mean. C. Quantification of the length of protrusions in three mSCLC cell lines (KP22, no visible protrusions, 16T and N2N1G with protrusions). The average cell size in these experiments was ∼8 μm. Each dot represents a cell. N>10 fields were quantified in one biological replicate. Mean +/− s.d. is shown, Mann-Whitney test. D. Representative still images from time-lapse videomicroscopy analysis of 16T SCLC cells showing the dynamic nature of the protrusions (from Movie S1). E. Quantification of the saltatory movements of three mSCLC cell lines as indicated. Note the correlation between the presence of protrusions and the ability of making longer steps (longer than the average cell size). Each dot represents a cell. N>10 fields were quantified in one biological replicate. Mean +/− s.d. is shown, Mann-Whitney test. F-H. Example of single cell movement over time for each of the three mSCLC cell lines.

We quantified the length of protrusions and found that they were often 5 to 10 times longer than the diameter of the cell body (∼8 µm) (Figure 2C). The length and the frequency of these axon-like protrusions suggested that they might influence the behavior of SCLC cells. We investigated and quantified the features of SCLC cells with and without protrusions using time-lapse microscopy. Initial observations of mouse SCLC cells showed that the protrusions were very dynamic (Figure 2C and Movie S1). In these movies, we noticed that the protrusions resembled cellular processes that have been described in neuroblasts and with the movement of SCLC cells along these protrusions reminiscent of neuroblast chain migration (Oudin *et al.*, 2011, Lois *et al.*, 1996, Zhou *et al.*, 2015). Indeed, when we quantified the movement of SCLC cell along protrusions, SCLC cell lines that form protrusions (16T and N2N1G cell lines) displayed increased saltatory activity compared to SCLC cells that do not form protrusions (KP22 cell line) (Figure 2D-H and Movies S2-4). The velocity of SCLC cells that form protrusions was also greatly increased compared to cells that do not form protrusions (Figure S2D).

Together, these results indicate that SCLC cells can generate axon-like protrusions and that these projections facilitate migration in a manner that is qualitatively similar to neuroblast migration during brain development.

### Loss of Axon-like protrusions inhibits the migration of SCLC cells

To investigate the functional importance of these axon-like protrusions, we focused on 13 genes (out of the 69 genes selected above) that encode for proteins involved in diverse aspects of axon formation, axon guidance, and neuronal migration (Table S4). These 13 genes are all expressed in at least a subset of human SCLC tumors (Figure S4A) (data from (George *et al.*, 2015)). We excluded gene families for which functional overlap and compensatory mechanism were likely. STRING analysis and literature searches confirmed that these 13 candidates had a significant connection with biological processes related to neurogenesis and the regulation of neuron projection development but that the proteins were not all directly connected and thus likely contribute to distinct aspects of these biological processes (Figure S4B and Table S5). In the 25 human SCLC cell lines analyzed in the Cancer Dependency Map project, knock-down of these 13 genes rarely affected the expansion of SCLC cells in culture, consistent with these genes influencing aspects of cell physiology not related to the cell cycle (Table S6 and Figure S4C-D).

We first knocked-down each of these genes with two shRNAs in a murine SCLC cell line derived from a lymph node metastasis (N2N1G). We confirmed stable knockdown by RT-qPCR (Table S7) and quantified the development of protrusions in the monolayer culture assay. Knock-down of 11 of the 13 genes significantly reduced the number of protrusions with at least one shRNA (Figure 3A-B and Figure S5A). The observation that the knock-down of multiple factors normally implicated at distinct steps of axonal growth reduces the development of the protrusions from SCLC cells further bolsters the notion that these protrusions are similar to neuronal axons. Knock-down of the many genes involved in axon formation, axonal guidance, and neuronal migration also reduced cell migration in the same assay (Figure 3B). Quantification of cell migration showed that inhibition of migration correlated with loss of the axon-like protrusions (Figure 3C-D). We validated the knock-down for two of the top candidates, *Gap43* and *Fez1* genes, by immunoblot for the corresponding proteins in N2N1G cells (Figure S5B-C). We further validated the effects of knocking down these two factors on the growth of protrusions and cell migration in a second SCLC cell line (16T; Figure 3E-J and Figure S5D-E).

**Figure 3:**
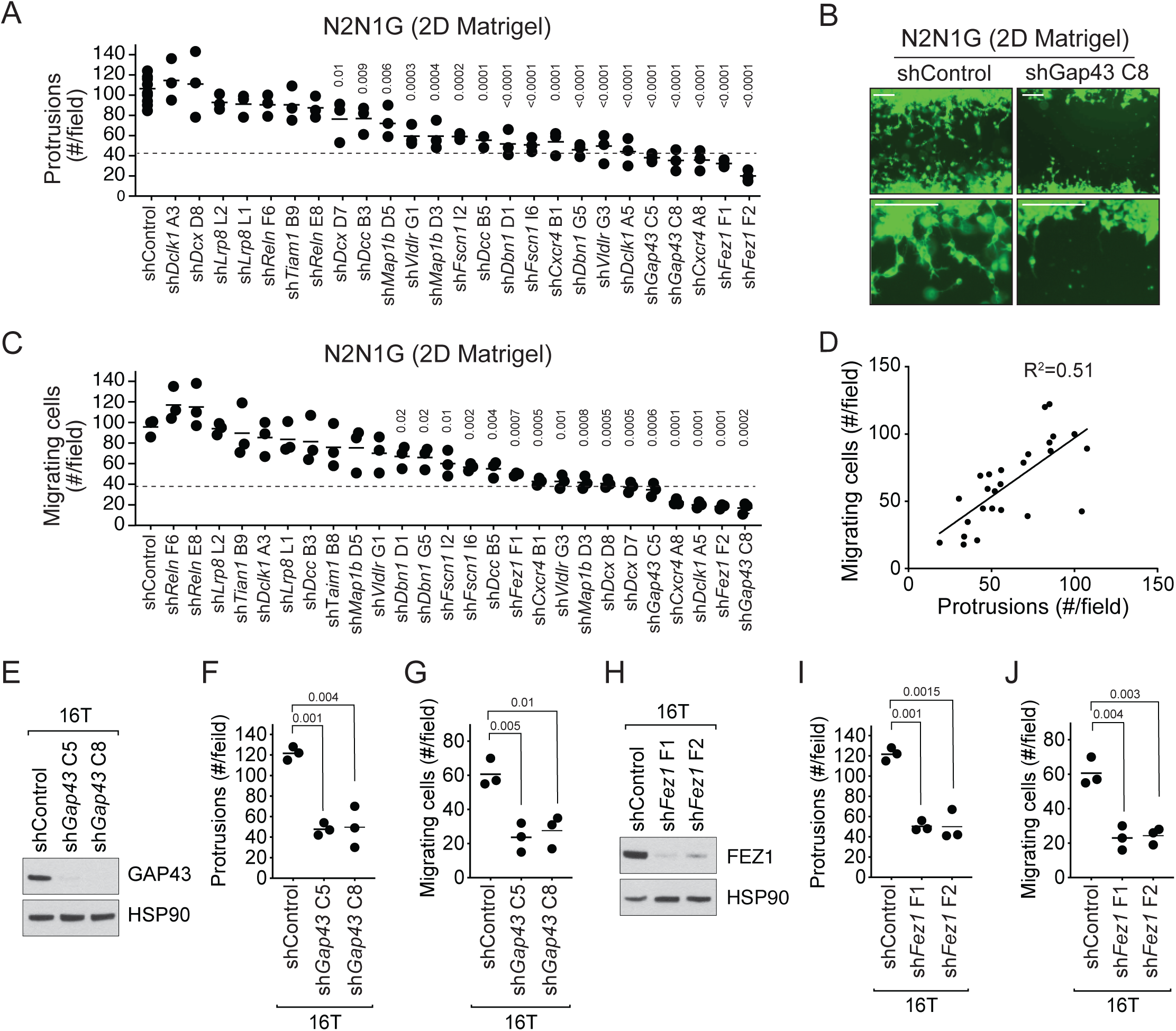
The axonal-like protrusions contribute to the migratory ability of SCLC cells in culture. A. Quantification of the number of cells with protrusions when mGFP-labeled N2N1G mSCLC cells were allowed to grow into a cell-free scratch generated in monolayer cultures under Matrigel. N=3 independent experiments (shControl, N=3 per experiment, total N=9 plotted together). An unpaired t-test was used for statistical analysis and p-values are shown. Only significant p-values are shown. The dotted line represents a 60% reduction compared to the mean value of the controls. B. Representative images of the data quantified in (A) and (C) with knock-down of *Gap43*. Scale bars, 100 μm. C. Quantification of the migration of cells with protrusions when mGFP-labeled N2N1G mSCLC cells were allowed to grow into a cell-free scratch generated in monolayer cultures under Matrigel. N=3 independent experiments. An unpaired t-test was used for statistical analysis and p-values are shown. Only significant p-values are shown. The dotted line represents a 60% reduction compared to the mean value of the controls. D. Correlation of the data in (A) and (C) using the mean value for each knock-down. Pearson correlation R^2^ value is shown. E and H. Immunoblot analysis of GAP43 or FEZ1 levels, respectively, in control and knock-down 16T mSCLC cells. HSP90 is a loading control. F and I. Quantification of the number of cells with protrusions as in (A) with 16T mSCLC cells and *Gap43* or *Fez1* knock-down, respectively (N=3). An unpaired t-test was used for statistical analysis and p-values are shown. G and J. Quantification of the migration of cells with protrusions as in (B) with 16T mSCLC cells and *Gap43* or *Fez1* knock-down, respectively (N=3). An unpaired t-test was used for statistical analysis and p-values are shown.

Together, these data show that SCLC cells with axon-like protrusions migrate in culture similar to what has been described for neuroblasts and that disruption of these protrusions by knocking-down a variety of genes involved in axonogenesis and neuronal migration also affects SCLC migration.

### Knock-down of genes associated with the formation of protrusions results in decreased metastatic potential

The link between axon-like protrusions and migration *in vitro* led us to investigate whether these axon-like protrusions promote the metastatic ability of SCLC cells *in vivo.* In support of this idea, we found that the expression of neuron-specific class III beta-tubulin and TAU was barely detectable in non-metastatic tumors in the lungs of *TKO* mice 3 months after cancer initiation while a majority of later stage tumors stained strongly positive for both proteins (Figure S6A-B).

To test the role of these protrusions in the metastatic process *in vivo*, we investigated whether SCLC cells with *Gap43* or *Fez1* knocked-down had reduced metastatic ability. The products of these genes are thought to regulate axonal development in entirely distinct manners but knock-down of each reduced the formation of protrusions and cell migration in culture. We first assessed whether *Gap43* and *Fez1* knock-down reduced the metastatic ability of mouse N2N1G SCLC cells after transplanting control and knock-down cells into recipient mice and assessing metastasis formation 4-5 weeks after intravenous injection. Knock-down of each of these pro-protrusion factors significantly reduced the number of metastases as assessed by tumor counts at the surface of the liver (Figure S7A-B). To determine whether GAP43 and FEZ1 are simply required for tumor growth *in vivo*, we transplanted *Gap43* and *Fez1* knock-down cells subcutaneously and quantified tumor growth. Knock-down of these genes had no effect on subcutaneous tumor growth suggesting that the effects on metastatic ability likely represent the disruption of phenotypes uniquely associated with the metastatic process (Figure S7C). We repeated these experiments with two independent shRNAs for each gene in both N2N1G and 16T SCLC cells, which confirmed that *Gap43* and *Fez1* knock-down inhibits the formation of liver metastases after intravenous injection of SCLC cells (Figure 4A-H and Figure S7D-E).

**Figure 4:**
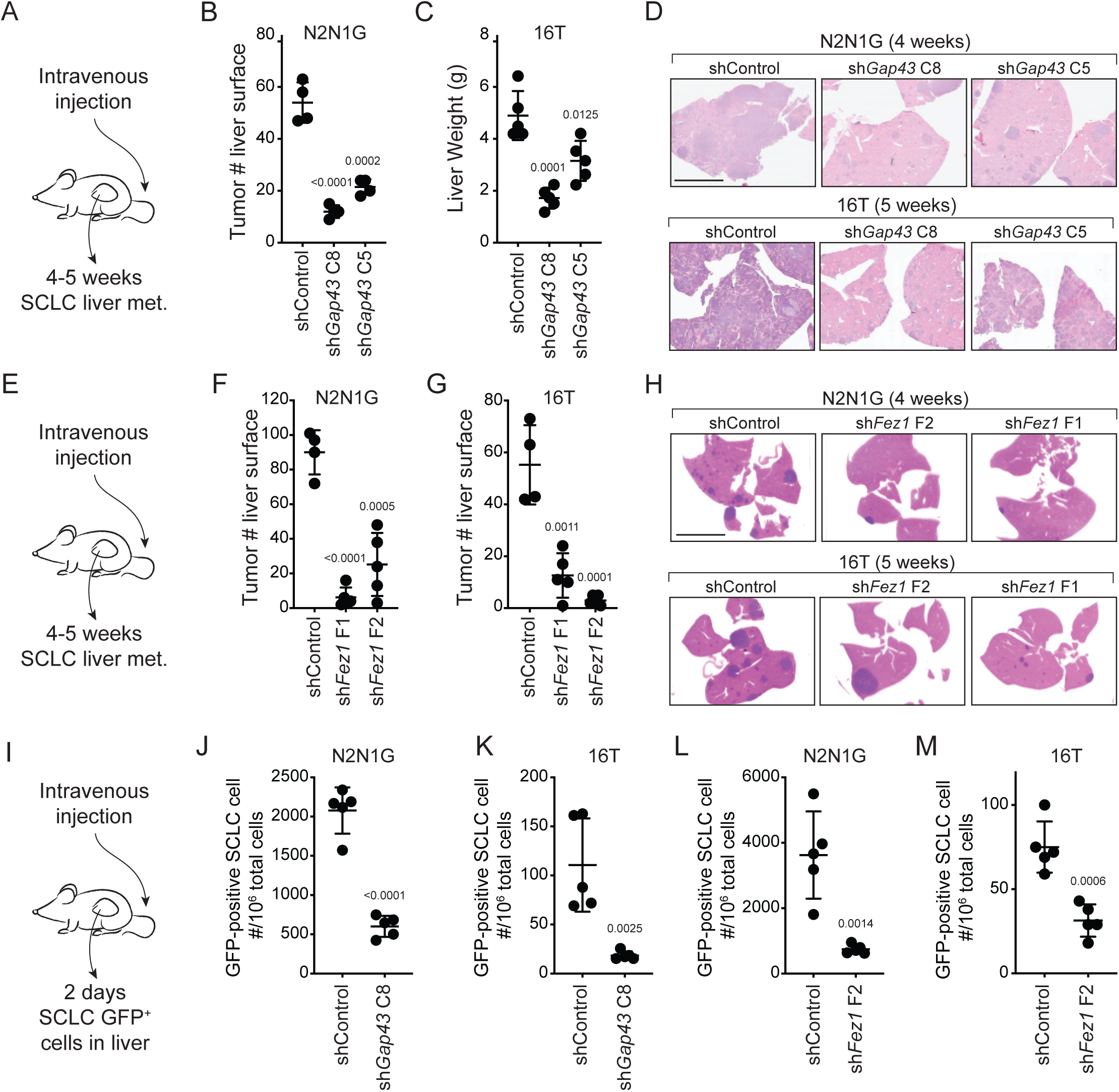
Genes involved in the generation of protrusions also control the formation of metastases. A. Diagram of the approach to investigate the formation of liver metastases (met.) after intravenous injection of SCLC cells. B-C. Quantification of the number of metastases 4 and 5 weeks after intravenous injection of N2N1G and 16T mSCLC cells, respectively, with control knock-down or knock-down of *Gap43* with two independent shRNAs. For N2N1G, tumors at the surface of the liver were quantified on the liver surface, as shown in Supplementary Figure S7D. Too many tumors were present with the 16T cell line and the control shRNA and quantification was thus performed by measuring liver weight. N=4-5 mice per condition in one biological replicate. Mean +/− s.d. unpaired t-test. D. Representative hematoxylin and eosin (H&E) images of liver sections of mice in (B-C). Scale bars, 5 mm. E-H. As shown in (A-D) for *Fez1* knock-down. See Supplementary Figure S7E for representative images with N2N1G cells for the quantification in (F-G) of tumors at the surface of the liver. N=4-5 mice per condition in one biological replicate. Mean +/− s.d. is shown, unpaired t-test. I. Diagram of the approach to investigate early steps in liver metastasis, 2 days after intravenous injection. J-M. Quantification of the number of GFP^positive^ (GFP^+^) N2N1G and 16T mSCLC cells 2 days after intravenous injection. See Supplementary Figure S7F-I for representative flow cytometry. N=5 mice per condition in one biological replicate. Mean +/− s.d., unpaired t-test.

The absence of growth defects in subcutaneous tumors following *Gap43* and *Fez1* knock-down suggested that these genes may affect earlier steps of the metastatic cascade. To test this, we performed similar intravenous transplant experiments but quantified the presence of SCLC cells in the liver 2 days after injection (Figure 4I). Quantification of GFP^positive^ cancer cells in the liver by flow cytometry documented a significant reduction in metastatic seeding by SCLC cells with *Gap43* and *Fez1* knocked-down (Figure 4J-M and Figure S7F-I). Thus, loss of genes associated with the formation of axon-like protrusions affects early metastatic seeding of SCLC cells in the liver, which ultimately translates to reduced metastatic burden.

## DISCUSSION

While metastasis remains a major cause of morbidity and mortality in SCLC patients, its underlying mechanisms remain poorly understood and no therapeutic strategies exist to prevent or target it. Here we investigated the function of neuronal gene expression programs in metastatic SCLC. We found that SCLC cells can grow axon-like protrusions and that these protrusions contribute to the migratory and metastatic phenotypes of these cells. This study identifies a cellular mechanism by which a neuroendocrine-to-neuronal transition promotes metastasis of SCLC cells.

The expression of neuronal factors in SCLC has been known for more than three decades and has been used as a marker for disease progression (Carney *et al.*, 1982, Cutz, 1982, Broers *et al.*, 1987, Anderson *et al.*, 1988). However, whether neuronal programs in SCLC cells play a direct role in SCLC progression had not been rigorously investigated. We uncovered the growth of axon-like protrusions as one functional aspect of neuronal differentiation in SCLC and provide data to support a role for these protrusions in migration and metastasis. It is likely that other phenotypes usually associated with neurons beyond these axon-like protrusions also contribute to the expansion and the spread of SCLC cells. Beyond facilitating metastatic seeding to the liver, these axon-like protrusions may have other functions, including helping SCLC cells migrate within the primary tumor, intravasate into the bloodstream, and move within the parenchyma during metastatic expansion (Shibue *et al.*, 2012). Future investigation of the roles of axon-like protrusions in SCLC will likely benefit from additional genetic analyses as well as high-resolution *in vivo* imaging methods. Recent evidence suggests that several other human tumor types also increase the expression of neuronal programs as they become more metastatic, especially to the brain (Wingrove *et al.*, 2019). It will be important for future studies to determine if aspects of the neuronal program also contribute to the striking ability of SCLC cells to seed and expand in the brain (Lukas *et al.*, 2017).

Our data indicate that SCLC metastasis is facilitated by the development of axon-like protrusions, but other molecular mechanisms certainly also increase the probability that a cancer cell will successfully overcome all the hurdles that limit the development of tissue destructive metastases. For instance, we found that knock-down of *Dcx* (coding for Doublecortin) has little to no effect on the number of protrusions but strongly inhibits migration in our 2D Matrigel assay (Figure 3A-B), thus suggesting that Doublecortin promotes SCLC migration independent from any impact of protrusion formation.

The formation of protrusions in SCLC cells is controlled by pathways previously implicated in the formation of axons and the migration of neuronal cells but it is unclear how the expression of these pro-protrusion genes is coordinated. We and others have identified a role for the NFIB transcription factor in SCLC metastasis and the induction of gene programs linked with axonogenesis and neuronal migration (Denny *et al.*, 2016, Semenova *et al.*, 2016, Wu *et al.*, 2016). However, overexpression of NFIB in naturally NFIB^low^ cell lines is not sufficient to induce the growth of protrusions in SCLC cells (unpublished observations). Thus, the upstream factors that control these neuronal programs in SCLC remain to be characterized. Accumulating evidence indicates the existence of several subtypes of SCLC, which are defined by the expression of key transcription factors (Rudin *et al.*, 2019). The murine cell lines used in this study are of the “SCLC-A” subtype (driven by the transcription factor ASCL1) but the human cell line NCI-H446 and the PDX model LU86 (Saunders *et al.*, 2015) belong to the “variant” subtype (SCLC-N, driven by the transcription factor NEUROD1). This suggests that the ability to grow protrusions may exist across subtypes. Possibly a combination of genetic and epigenetic factors contributes to the ability of SCLC to grow protrusions. Adhesion molecules and other factors in the tumor microenvironment are also likely to contribute to the formation of protrusions *in vivo* (Guo *et al.*, 2000).

Could an understanding of the molecular and cellular processes related to axon-like protrusions in SCLC cells ultimately be translated into clinical benefit for SCLC patients? Previous studies on SCLC have targeted the CXCR4 chemokine receptor due to its role in cell adhesion and migration and its expression in SCLC cells (Burger *et al.*, 2003, Teicher, 2014, Taromi *et al.*, 2016). CXCR4 also contributes to the formation of axon-like protrusions (Figure 3). In a recent clinical trial in SCLC patients, CXCR4 inhibition was well tolerated but this inhibition did not significantly reduce disease progression (Salgia *et al.*, 2017). Mechanisms that drive the ability of cancer cells to overcome early barriers of metastatic seeding will likely need to be employed in specific settings where inhibition of the metastatic process would logically provide clinical benefit. For example, in patients with resectable SCLC, inhibition of pro-metastatic pathways in the neo-adjuvant and/or adjuvant setting could reduce the frequency or multiplicity of metastatic relapse.

More generally, the transition from a neuroendocrine state to a state where neuroendocrine differentiation is decreased but neuronal differentiation is increased may be related to the exceptional plasticity of SCLC cells (reviewed in (Yuan *et al.*, 2019)). Epithelial-to-mesenchymal transition (EMT) is thought to contribute to migration, metastasis, and resistance to treatment in many cancer contexts and may play a role in SCLC (Allison Stewart *et al.*, 2017, Krohn *et al.*, 2014, Canadas *et al.*, 2014, O’Brien-Ball and Biddle, 2017, Singh and Settleman, 2010). Vascular mimicry (or epithelial-to-endothelial transition (EET) (Yuan *et al.*, 2019)) may also contribute to tumor growth and response to treatment in SCLC (Williamson *et al.*, 2016). Similarly, Notch-induced dedifferentiation to a non-neuroendocrine state can generate an intra-tumoral niche that protects neuroendocrine SCLC cells (Lim *et al.*, 2017). Based on our results and recent observations in other cancers (Wingrove *et al.*, 2019), we propose that an epithelial-to-neuronal transition contributes to key aspects of cancer metastasis. Further characterization of this neuronal state in both neuroendocrine and non-neuroendocrine cancers is likely to uncover novel mechanisms of cancer progression and may ultimately offer new insight into metastasis-blocking strategies in the clinic.

## MATERIAL AND METHODS

### Mouse model

All experiments were performed in accordance with Stanford University Institutional Animal Care and Use Committee guidelines. *Trp53*^*flox*^, *Rb1*^*flox*^, *p130*^*flox*^, and *R26*^*mTmG*^ mice have been described (Denny *et al.*, 2016, Muzumdar *et al.*, 2007). Tumors were initiated by inhalation of Adeno-CMV-Cre (University of Iowa Vector Core, Iowa city, Iowa) as described in (Denny *et al.*, 2016), following a published protocol (DuPage *et al.*, 2009).

### Cell culture

All murine and human SCLC cell lines used in this study grow as floating aggregates and were cultured in RPMI with 10% FBS, 1×GlutaMax, and 100 U/mL penicillin-streptomycin (Gibco, Thermo Fisher Scientific, Waltham, MA). Human cell lines were originally purchased from ATCC and cell identities were validated by Genetica DNA Laboratories using STR analysis. NJH29 SCLC cells were derived from a patient-derived xenograft (PDX), which has been described (Jahchan *et al.*, 2013). The LU86 and LU102 models were obtained from Stemcentrx (Saunders *et al.*, 2015). The JHU-LX102 (LX102) model was a kind gift from Dr. Watkins (Leong *et al.*, 2014). The murine cell lines were described (Denny *et al.*, 2016, Yang *et al.*, 2018). Briefly, 16T and KP22 cells are from individual primary tumors from the lungs of *Rb/p53 DKO* mice. N2N1G cells were derived from a lymph node metastasis in an *Rb/p53/p130 TKO; Rosa26*^*mTmG*^ mouse. 6PF cells were derived from metastatic cells in the plural fluid in an *Rb/p53/p130 TKO; Rosa26*^*mTmG*^ mouse. All cell lines were confirmed to be mycoplasma-negative (MycoAlert Detection Kit, Lonza, Basel, Switzerland).

### *In vitro* 2D Matrigel migration and protrusion assay

Silicone inserts (ibidi 80209, Grafelfing, Germany) were attached to wells in 12-well (up to two inserts) or 24-well (one insert) plates pre-coated with poly-D-lysine for 15 minutes (Sigma-Aldrich, St. Louis, MO). ∼8×10^5^ cells were seeded to each chamber of the insert in 100 µL resulting in cells at ∼80-90% confluency. After at least 6 hours, the inserts were carefully removed and 0.75-1 mL of a 1:1 Matrigel (Corning, Corning, NY)-cell culture media mix was slowly added to cover each well. 1 mL of cell culture media was added on top of the solidified Matrigel to prevent drying. For quantification of cell migration and protrusions, the number of cells and the number of protrusions were counted in the gap at 10x under the microscope. The time points (between 36 hours and 96 hours) were dependent on the growth rate of the cell populations.

### Live imaging of cell migration and quantification

SCLC cells were plated as described in the 2D Matrigel migration assay and cultured for 24 hours before imaging. Then 10x DIC images were collected every 15 minutes for 25 hours using a Zeiss LSM 710 confocal microscope (Zeiss, Oberkochen, Germany) with a live imaging chamber set to 37°C, 5% CO2. To quantify the time-lapse movies, we examined nuclear movement and process length (as described in (Oudin *et al.*, 2011)) using the FIJI software (NIH, Bethesda, USA). The position of the cell nucleus was tracked in each frame using the Manual Tracking plugin to obtain the distance migrated by the nucleus per frame and the average cell velocity over the entire movie. Neuronal cell migration occurs via three steps: the cell extends a leading process, the nucleus translocates into the leading process via nucleokinesis, and the cell loses its trailing process. To quantify translocation events, we quantified the fractions of steps taken by each that were over 8um, which represents the length of one cell body and a nuclear translocation event. The process length was calculated by tracing a line from the cell body to the tip of the leading process about 6hrs into the movie. Over 30 cells were tracked and analyzed per condition.

### Immunostaining of cells in cultures

Cells were fixed with 4% PFA for 15 minutes, permeabilized with 0.1% Triton and stained for Tuj1 (1:500, BioLegend 801213, San Diego, CA), TAU (1:1000, Dako A0024, Santa Clara, CA), and MAP2 (1:500, EMD Millipore AB5622, Burlington, MA), and with a goat anti-rabbit secondary antibody (Invitrogen). Membrane GFP was stained (Abcam ab13970, Cambridge, UK) to mark SCLC cells and the expression of the other neuronal markers were checked using a fluorescence scope (Zeiss LSM 880). Staining was quantified by counting directly under the microscope (at 40x magnification).

### Whole mount immunofluorescence staining and imaging of tumors

Detailed methods for whole mount immunofluorescence staining have been described (Yang *et al.*, 2018). Subcutaneous tumors with 5-10% GFP^positive^ labeled cells mixed with non-GFP labeled SCLC tumor cells were dissected and were fixed in 4% paraformaldehyde and sectioned with a vibrating blade microtome at 500 µm thickness. Tumor slices were optically cleared using the CUBIC method, comprised of a three-hour incubation at room temperature in CUBIC 1 reagent and long-term storage in CUBIC 2 at 4°C (Susaki *et al.*, 2015). Sections were imaged using a Zeiss LSM 780 laser scanning confocal microscope.

For DiI staining and imaging, subcutaneously transplanted human SCLC xenograft were harvested after 3 weeks of growth and cut into 500mm ∼1 cm thick slices. Tumor pieces were stained with the red fluorescent tracer DiI (D282, Thermo Fisher Scientific) in a spot-wise manner, incubated in 37°C, 5% CO_2_ chamber for 20min and washed 3 times with PBS+10%FBS to remove excess DiI before imaging. Images were collected using a Leica SP5 scope (Leica, Buffalo Grove, IL) with a water immersion lens.

### Histology and immunohistochemistry

Mouse tumor samples were fixed in 4% formalin and paraffin embedded. Hematoxylin and Eosin (H&E) staining was performed using standard methods. For immunohistochemistry, we used antibodies to GFP (Abcam ab6673), UCHL1 (Sigma-Aldrich HPA005993), Tuj1 (BioLegend 801213), and TAU (Dako A0024).

Tissue microarrays (LC818a, US Biomax, Rockville, MD) were stained for TAU and scored by a board-certified pathologist on a three point scale as follows: 0 = negative or weak staining of less than 10% cells, 1 = moderate intensity staining, 2 = strong intensity staining.

### Candidate gene knockdown

Stable knockdown of candidate genes was performed using lentiviral pLKO vectors and puromycin-resistance selection (Sigma-Aldrich). For lentivirus production, 7.5×10^6^ HEK293T cells were seeded into 10 cm dishes and transfected with the vector of interest using PEI (Polysciences 23966-2, Warrington, PA) along with pCMV-VSV-G (Addgene #8454) envelope plasmid and pCMV-dR8.2 dvpr (Addgene #8455) packaging plasmid. The medium was changed 24 hours later. Supernatants were collected at 36 hours and 48 hours, passed through a 40 µm filter and applied at full concentration to target cells. Two days after transduction cells were selected with Puromycin (2 µg/mL, Thermo Fisher Scientific, Waltham, MA) for at least 1 week. Knockdown was confirmed by RT-qPCR as in (Denny *et al.*, 2016) and immunoblot analysis. Table S7 shows the sequences of the oligonucleotides used to knock down the candidate genes. Note that the expression of the shRNAs targeting GFP partially decreased GFP expression, but cancer cells were still GFP^positive^ and could be well-detected by flow cytometry.

### Immunoblot analysis

GAP43 (Abcam), FEZ1 (Cell Signaling, Danvers, MA), and HSP90 (BD Transduction Laboratories, San Jose, CA) antibodies were used to confirm the knockdown of each gene at the protein level. Briefly, denatured protein samples were run on 4-12% Bis-Tris gels (NuPage, Thermo Fisher Scientific, Waltham, MA) and transferred onto PVDF membrane. Primary antibody incubations were followed by secondary HRP-conjugated anti-mouse (Santa Cruz Biotechnology, Santa Cruz, CA) and anti-rabbit (Santa Cruz Biotechnology) antibodies and membranes were developed with the ECL2 Western Blotting Substrate (Pierce Protein Biology, Thermo Fisher Scientific).

### Transplantation assays

For long-term metastasis assays, 3×10^4^ of N2N1G cells or 1×10^5^ of 16T cells were injected intravenously injected into the lateral tail vein of NOD.*Cg-Prkdc*^*scid*^*Il2rg*^*tm1Wjl*^/SzJ (NSG) mice (The Jackson Laboratories, Bar Harbor, ME – Stock number 005557). Mouse livers were harvested at 4-6 weeks after injection. Tumor number was quantified by directly counting on liver surface and also quantified by counting tumor number or areas on the H&E sections. For subcutaneous injection, 5×10^4^ cells were resuspended in 100 µL PBS and mixed with 100 µL Matrigel (Corning, 356231, Corning, NY) with 4 injection sites per mouse. For both subcutaneous and intravenous injections, SCLC cells were transplanted into age-matched gender-matched NSG mice. For short-term tumor seeding assays, 2×10^7^ of N2N1G cells or 5×10^7^ of 16T cells were transplanted intravenously into the lateral tail vein of NSG mice. N2N1G, derived from *Rb/p53/p130 TKO; Rosa26*^*mTmG*^ mouse, has endogenous GFP expression and 16T, derived from *Rb/p53 TKO* mouse, was stained by live cell stain CFSE (Thermo Fisher Scientific, C34554) and washed before intravenous injection. 2 days after transplantation, mouse livers were harvested, digested into single cell suspension and analyzed by FACS for the percentage of GFP^positive^ cancer cells. FACS data were analyzed by FlowJo.

### Pathway and process enrichment analysis

Metascape (metascape.org) was used to analyze the lists of genes involved in axonogenesis and neuronal migration. Metascape integrates data from KEGG Pathway, GO Biological Processes, Reactome Gene Sets, Canonical Pathways and CORUM (Zhou *et al.*, 2019). The analysis of interactions between the top 13 candidate genes was performed using STRING (string-db.org) (Szklarczyk *et al.*, 2018). The analysis of dependency upon knock-down was performed using the in the Cancer Dependency Map project (depmap.org/portal/) in February 2019 with the Combined RNAi (Broad, Novartis, Marcotte) data (Tsherniak *et al.*, 2017).

### Statistics

Statistical significance was assayed with GraphPad Prism software. The statistical tests used, the numerical p-values, and the number of independent replicates is indicated in the figure legends.

## ACKNOWLEDGEMENTS

We thank Pauline Chu and Jon Mulholland for technical assistance; Alexandra Orantes and Alyssa Ray for administrative support; Kang Shen, Pengpeng Li, Gregor Bieri, Nick Kramer, Gregory Giannone, and Olivier Rossier for helpful discussions; and members of the Winslow and Sage laboratories for helpful comments. We thank Dr. Charlie Rudin and Stemcentrx for the PDX models. We thank the Stanford Shared FACS Facility and Cell Sciences Imaging Facility. This work was supported by NIH R01 CA206540 (to JS) and in part by the Stanford Cancer Institute support grant (NIH P30 CA124435). DY was supported by a Stanford Graduate Fellowship and by a TRDRP Dissertation Award (24DT-0001). FQ was supported by a Damon Runyon Postdoctoral Fellowship. HC was supported by a Tobacco-Related Disease Research Program Postdoctoral Fellowship. C-HC was supported by an American Lung Association Fellowship. B.M.G. was supported by the Pancreatic Cancer Action Network – AACR Fellowship in memory of Samuel Stroum (14-40-25-GRUE), was a Hope Funds for Cancer Research Fellow supported by the Hope Funds for Cancer Research (HFCR-15-06-07), and is a recipient of an Emmy Noether Award from the German Research Foundation (DFG). MJO was supported by NIH R00 CA207866. J.S is the Harriet and Mary Zelencik Scientist in Children’s Cancer and Blood Diseases.

---

**Figure S1 (related to Figure 1).**
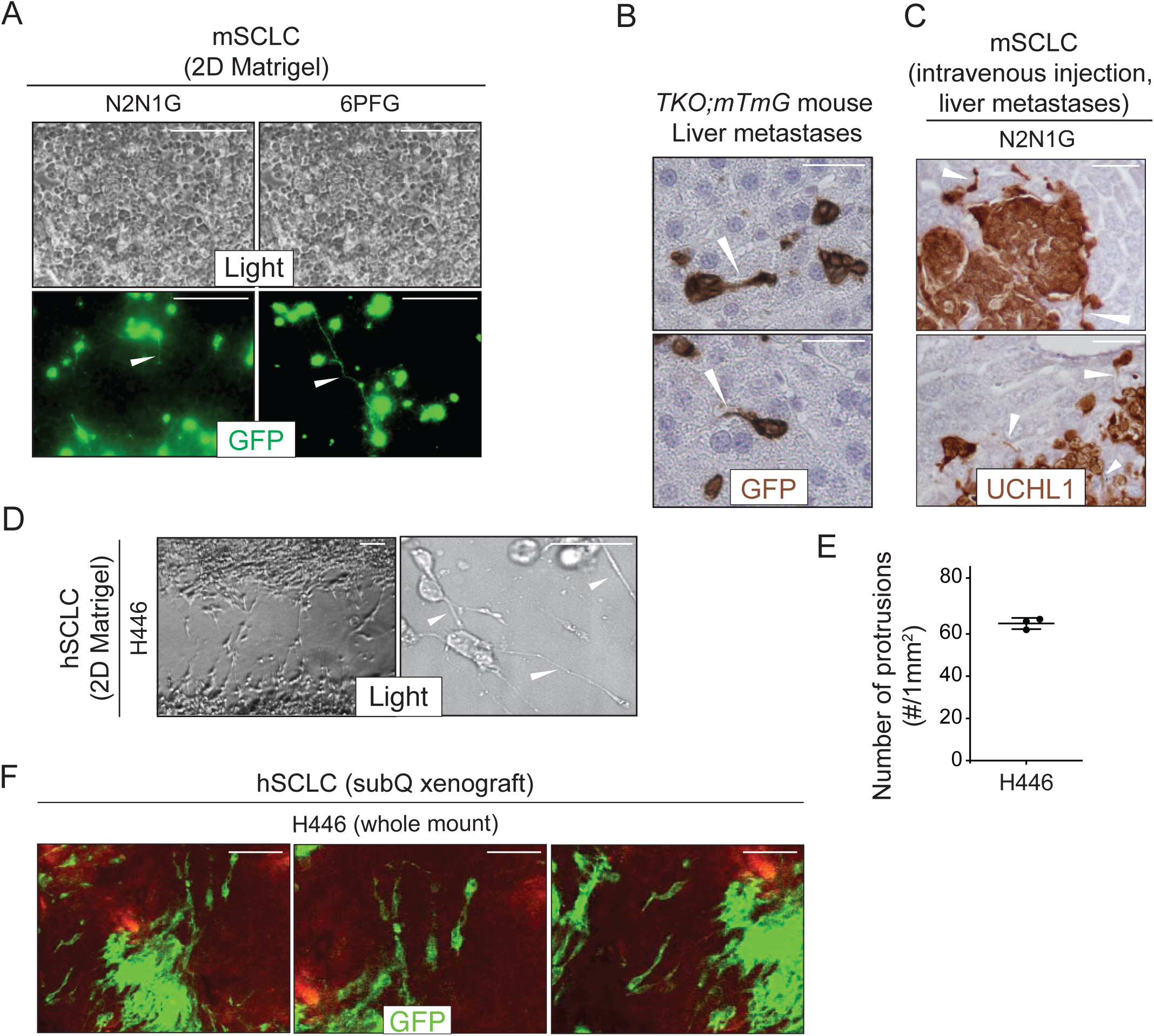
SCLC cells grow protrusions in culture and *in vivo*. A. Representative images of mSCLC N2N1G and 6PFG cells growing in dense culture from N=3 independent experiments. At the time of plating, 3-5% cells expressing membrane-GFP (mGFP, green fluorescence) were mixed and co-cultured with 95-97% SCLC cells that do not expressing GFP. Examples of protrusions are shown with white arrowheads. Scale bars, 100 μm. B. Representative images of mSCLC cells in the liver from the autochthonous *TKO;mTmG* model from N= 2 mice. Images were taken from micro-metastases. Immunostaining for GFP generates a brown signal. Protrusions are shown with white arrowheads. Hematoxylin (blue) stains the nucleus of cells. Scale bar, 20 μm. C. Representative images of liver sections from mice after intravenous injection of mSCLC N2N1G cells from N=3 mice. Immunostaining for the neuroendocrine marker UCHL1 (brown) outlines the shape of cells. Protrusions are shown with white arrowheads. Scale bars, 50 μm. D. Representative bright field images of human SCLC (hSCLC) NCI-H446 cells when cells are allowed to grow into a cell-free scratch generated in monolayer cultures under Matrigel. Protrusions are shown with white arrowheads. Scale bars, 40 μm. E. Quantification of (D). N=3 independent experiments. Mean +/− s.d. is shown. F. Representative whole mount images of hSCLC NCI-H446 cells growing as a subcutaneous tumor from N=4 independent xenografts from one experiment. At the time of injection, 10% of the SCLC H446 cells expressing membrane-GFP (mGFP) were mixed with 90% SCLC H446 cells not expressing GFP. Scale bars, 100 μm.

**Figure S2 (related to Figure 2).**
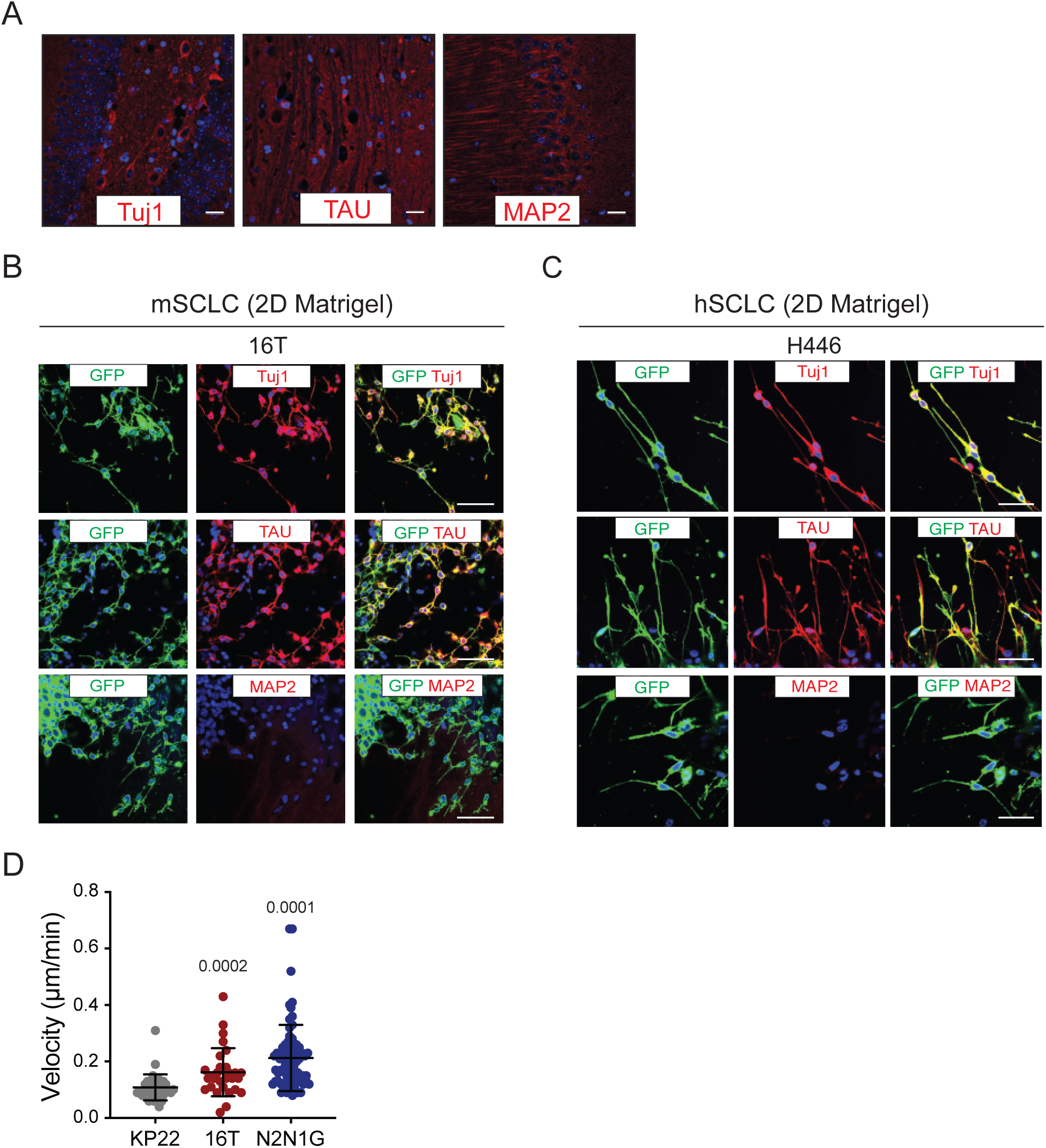
SCLC protrusions resemble axons and enable rapid cell movement. A. Representative fluorescence images of a mouse brain section stained with Tuj1, TAU, and MAP2 antibodies (positive controls, red). DAPI marks the nuclei of cells in blue. Scale bars, 50 μm. B-C. Representative fluorescence images of 16T mSCLC cells (B) and NCI-H446 hSCLC cells (C) expressing membrane-GFP (mGFP) and stained (red) for expression of the neuronal marker Tuj1, the axonal marker TAU, or the dentritic marker MAP2. DAPI marks the nucleus of cells in blue. Quantification is shown in Figure 2B. Scale bars, 50 μm. D. Quantification of velocity of mSCLC cancer cells from the three mouse SCLC cell lines indicated. Each dot represents a cell. Mean +/− s.d. is shown, Mann-Whitney test.

**Figure S3 (related to Figure 2).**
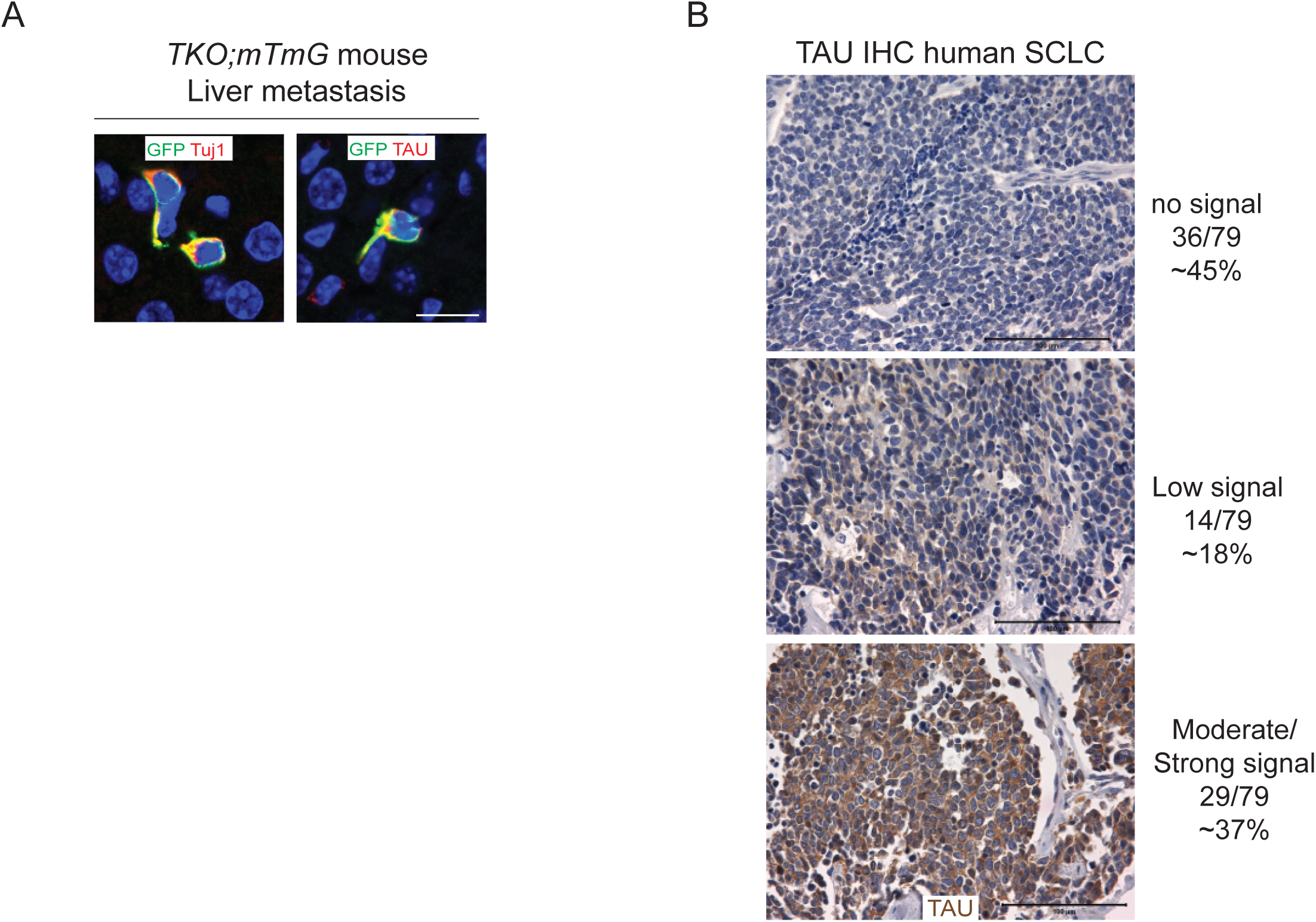
Mouse and human SCLC cells express axonal markers *in vivo*. A. Representative immunofluorescence staining of SCLC cells in the liver of a *TKO;mTmG* mouse (in which SCLC cancer cells express membrane GFP (GFP)). These cancer cells have protrusions positive for TAU and Tuj1. Images represent a merge of the GFP signal (green) and the signal for the TAU or Tuj1 antibodies (red). The nucleus of cells is labeled in blue by DAPI. Scale bar, 20 μm. B. Representative images of immunohistochemistry (IHC) for TAU (brown) on human SCLC tissue microarrays (N=79 human samples analyzed). The signal was evaluated by a certified pathologist (K.C.). Scale bars, 100 μm.

**Figure S4 (related to Figure 3).**
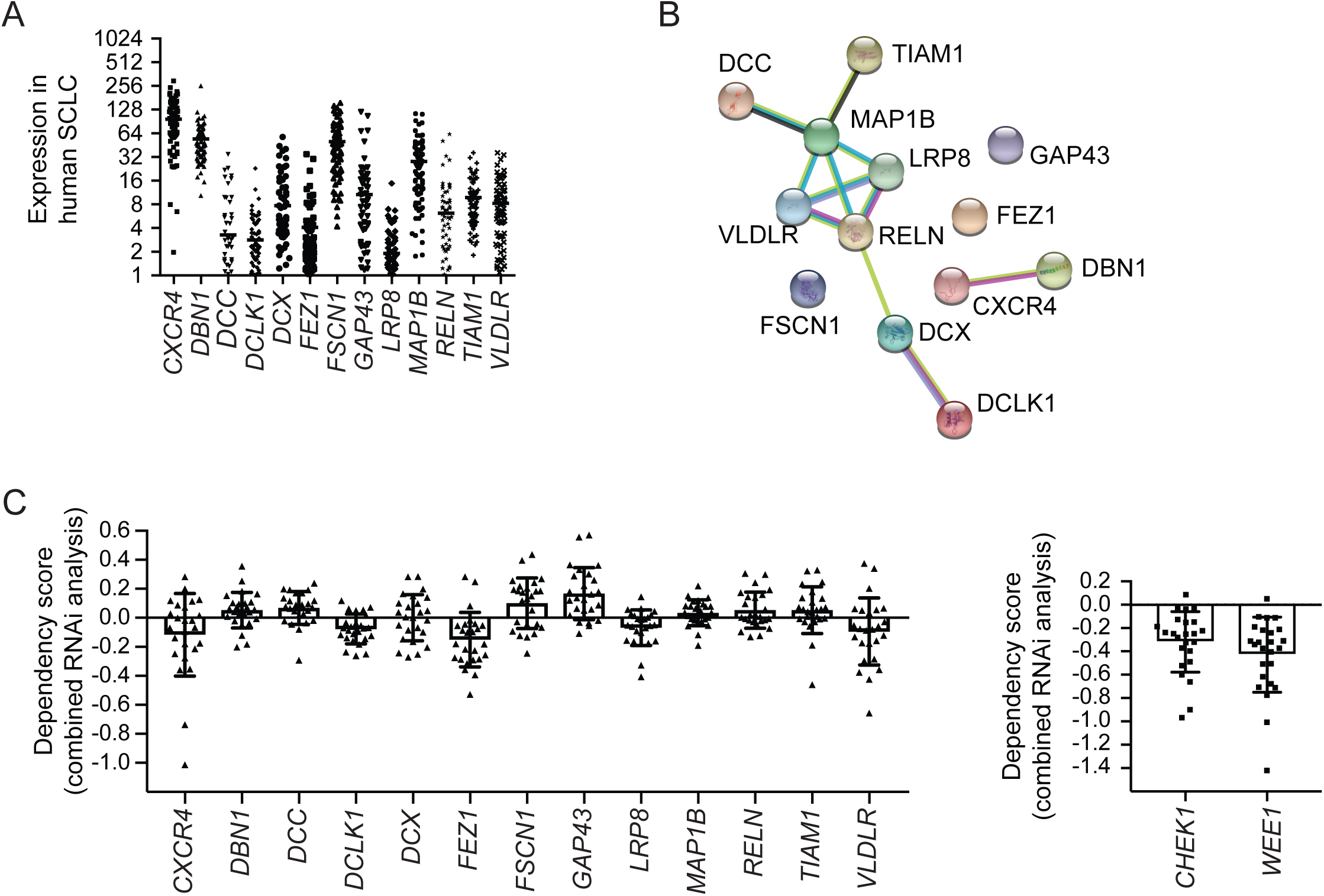
The 13 genes selected for their possible role in the formation of protrusions are expressed in human SCLC but do not play a key role in the expansion of SCLC cell populations. A. mRNA levels of candidate genes in human primary SCLC tumors (RNA-seq from George, Lim *et al.*, Nature, 2015). B. Network representation of the 13 candidates. Edges in the STRING analysis represent protein-protein associations but do not necessarily mean that they physically bind to each other. Blue edges represent known interactions from curated databases. Pink edges represent known experimentally-validated interactions. Others are predicted interactions, including text mining and co-expression (see string-db.org). C. DepMap analysis (depmap.org) of the requirement for the 13 candidate genes in 25 human SCLC cell lines (RNAi combined analysis). Note that in a number of cell lines, the knock-down of candidate genes results in a positive score, indicative of a better expansion upon knock-down. Even in cases where the scores are negative, the negative values are small (the data for the genes coding for the CHK1 and WEE1 kinases, which are considered therapeutic targets in SCLC, are shown on the right hand side).

**Figure S5 (related to Figure 3).**
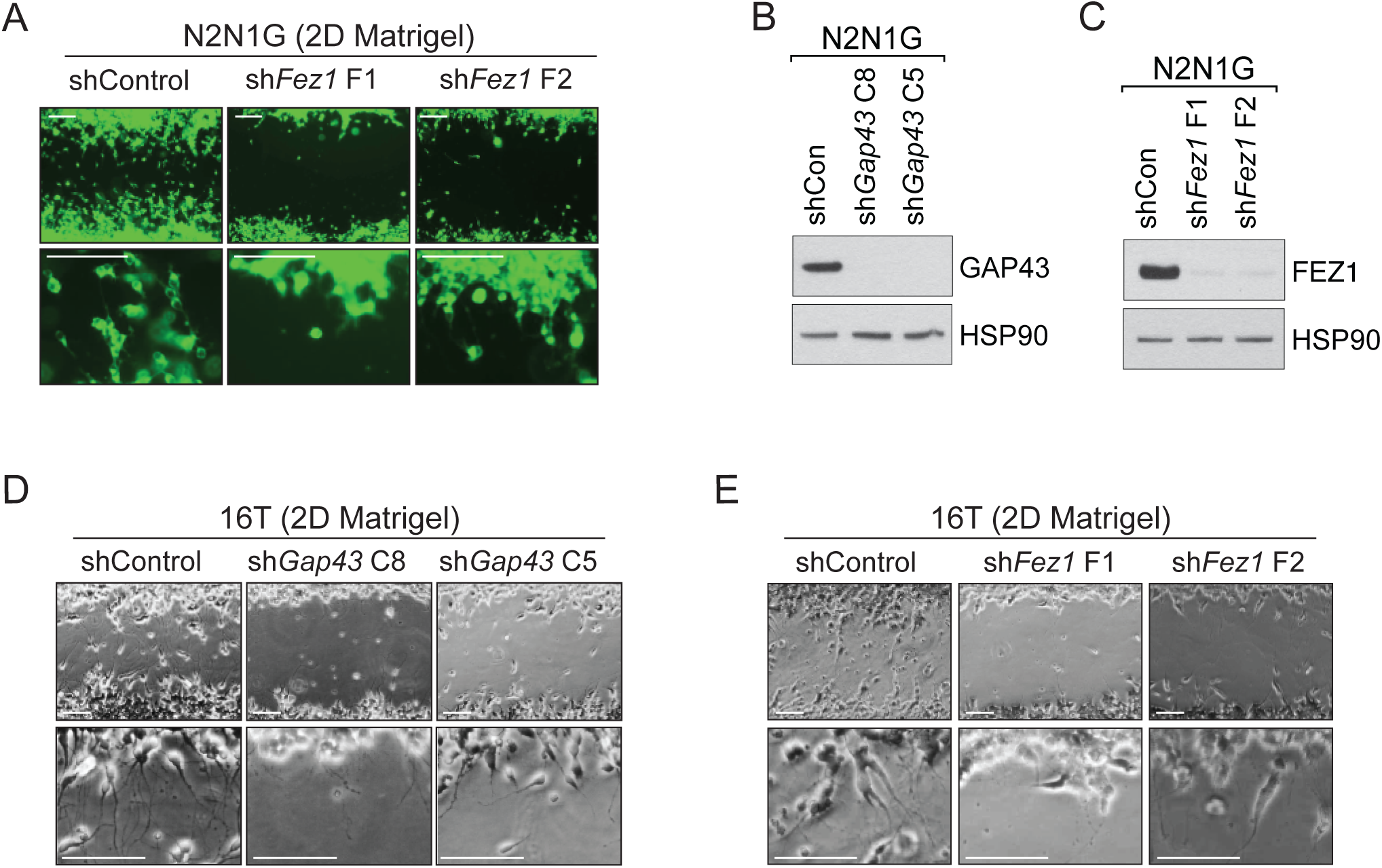
Knock-down of GAP43 and FEZ1 disrupts the formation of protrusions and cell migration in mouse SCLC cell lines in culture. A. Representative images of the data qsuantified in Figure 3A and 3C with knock-down of *Fez1*. Scale bars, 100 μm. B-C. Immunoblot analysis of GAP43 (B) or FEZ1 (C) levels in control and knock-down N2N1G mSCLC cells. HSP90 is a loading control. D-E. Representative images of the data with knock-down of *Gap43 (D)* or *Fez1* (E) in 16T cells. These data are quantified in Figure 3F-G (for GAP43) and Figure 3I-J (for FEZ1). The shControl targets GFP. Scale bars, 100 μm.

**Figure S6 (related to Figure 4).**
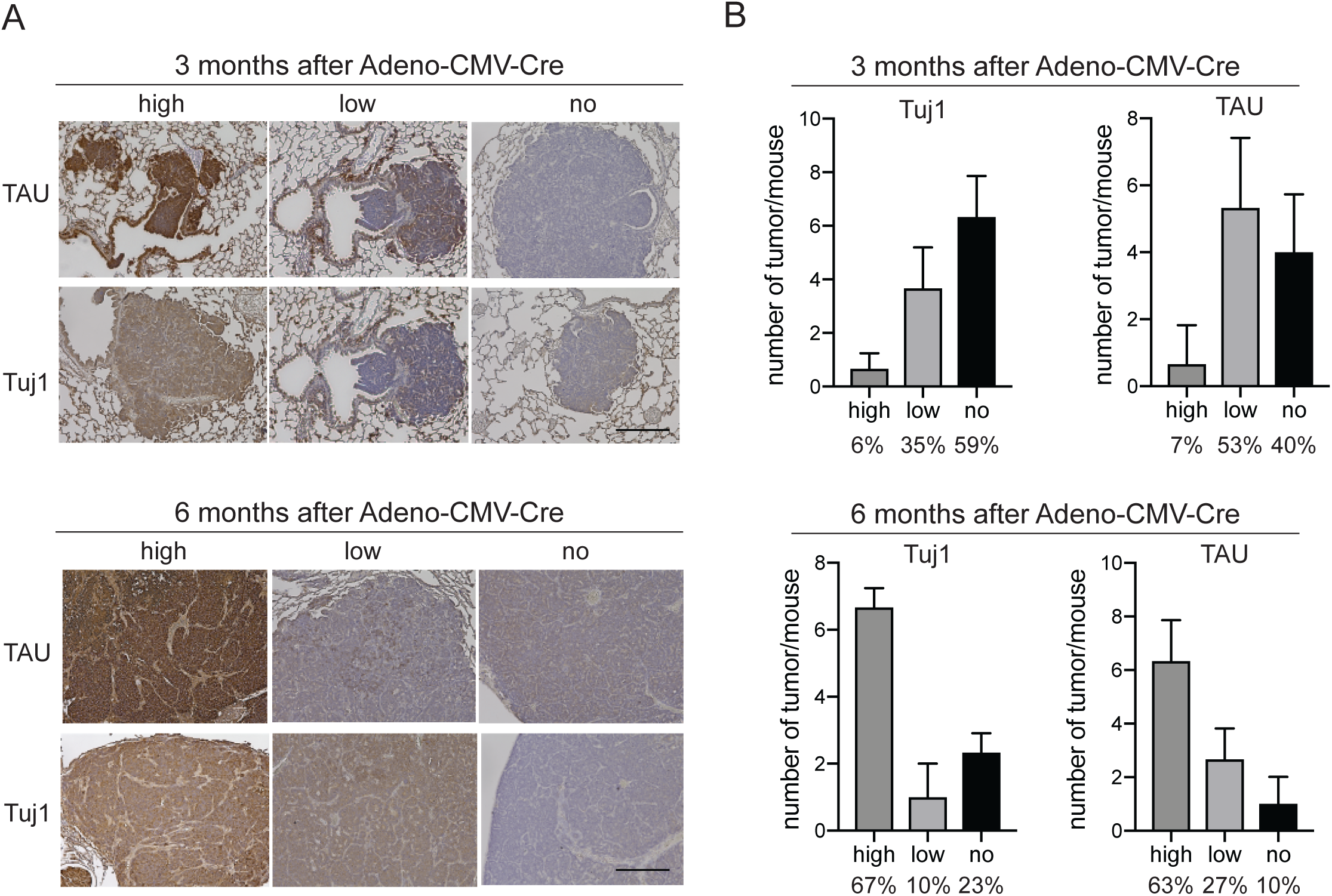
Increased expression of the axonal marker TAU in metastatic SCLC in the *TKO* mouse model. A. Representative images of immunohistochemistry experiments on lung sections from *TKO* mice 3 months and 6 months after SCLC initiation with Ad-CMV-Cre. None of the mice had metastases at the 3-month time point while all the mice analyzed had evidence of metastasis at the 6-month time point. The Tuj1 antibody marks neuronal tubulin and TAU is a marker of axons. Hematoxylin was used as a counterstain (purple). Scale bar, 100 μm. B. Quantification of (A), with N=30-32 tumors analyzed from N=3 mice at the 3-month time point and N=30 tumors analyzed from N=3 mice at the 6-month time point. Percentages are indicated.

**Figure S7 (related to Figure 4).**
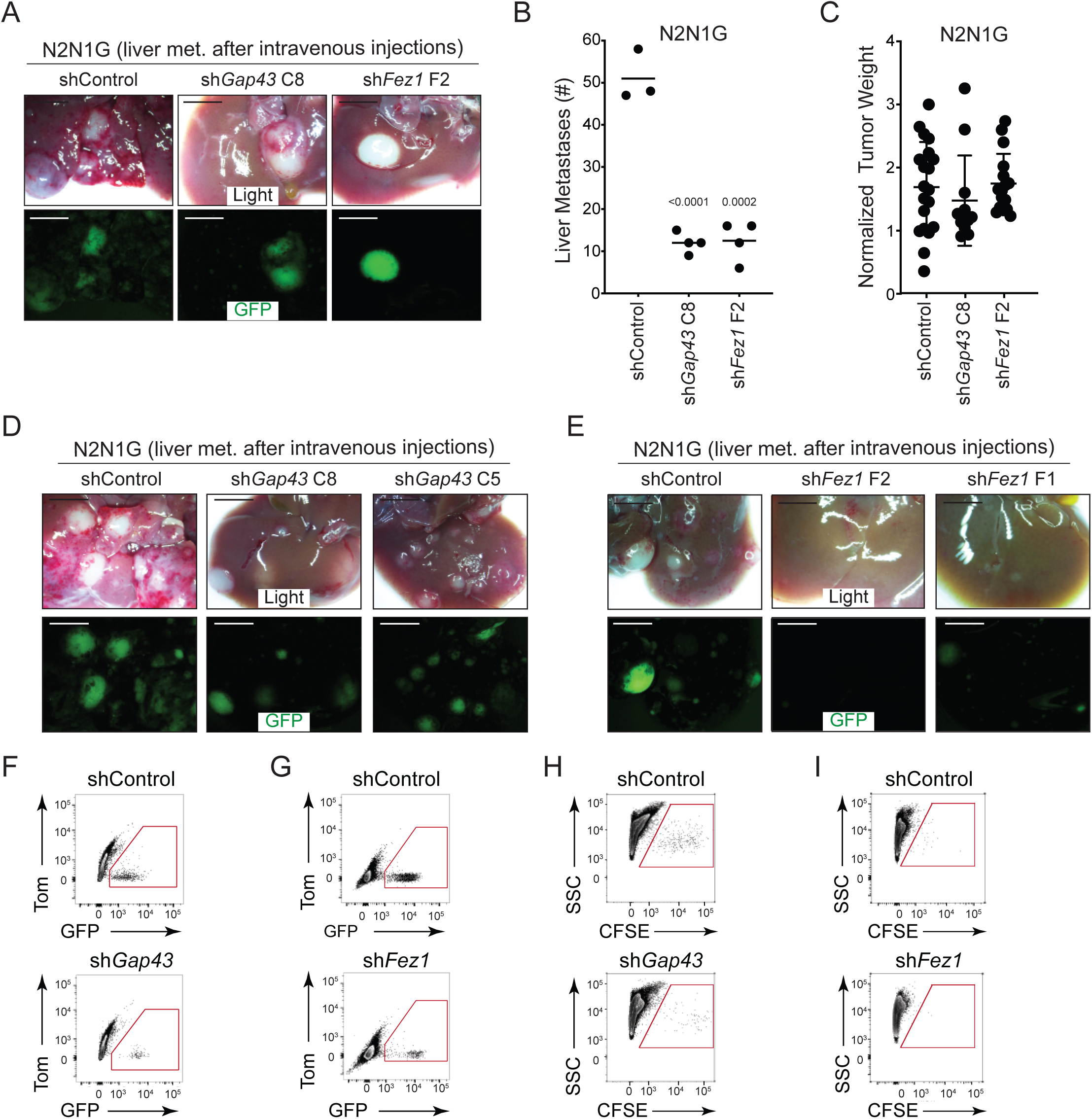
Reduced formation of metastasis upon knock-down of GAP43 and FEZ1 in SCLC cells. A. Representative live and epifluorescence images (GFP, green) of liver section of mice 4 weeks after intravenous injection of GFP-positive N2N1G mSCLC cells, with control knock-down or knock-down of *Gap43 or Fez1*. Scale bars, 5 mm. B. Quantification of (A). The bar is the mean, unpaired t-test. C. Quantification of tumor weight after subcutaneous injection of control and knock-down N2N1G cells. Values are not statistically significant by t-test. D-E. Representative bright light and epifluorescence images (GFP, green) of liver section of mice 4 weeks after intravenous injection of GFP-positive N2N1G mSCLC cells, with control knock-down or knock-down of *Gap43 or Fez1*. Scale bars, 5 mm. F-G. Representative flow cytometry quantification of GFP-positive N2N1G cells in the liver 2 days after intravenous injection. H-I. Representative flow cytometry quantification of CFSE-labeled 16T cells in the liver 2 days after intravenous injection.

